# Structural insights into the ion selectivity of the MgtE channel for Mg^2+^ over Ca^2+^

**DOI:** 10.1101/2021.12.29.474488

**Authors:** Xinyu Teng, Danqi Sheng, Jin Wang, Ye Yu, Motoyuki Hattori

## Abstract

MgtE is a Mg^2+^-selective ion channel whose orthologs are widely distributed from prokaryotes to eukaryotes, including humans, and play an important role in the maintenance of cellular Mg^2+^ homeostasis. Previous functional analyses showed that MgtE transports divalent cations with high selectivity for Mg^2+^ over Ca^2+^. Whereas the high-resolution structure determination of the MgtE transmembrane (TM) domain in complex with Mg^2+^ ions revealed a Mg^2+^ recognition mechanism of MgtE, the previous Ca^2+^-bound structure of the MgtE TM domain was determined only at moderate resolution (3.2 Å resolution), which was insufficient to visualize the water molecules coordinated to Ca^2+^ ions. Thus, the structural basis of the ion selectivity of MgtE for Mg^2+^ over Ca^2+^ has remained unclear. Here, we showed that the metal-binding site of the MgtE TM domain binds to Mg^2+^ ∼500-fold more strongly than Ca^2+^. We then determined the crystal structure of the MgtE TM domain in complex with Ca^2+^ ions at a higher resolution (2.5 Å resolution), allowing us to reveal hexahydrated Ca^2+^, which is similarly observed in the previously determined Mg^2+^-bound structure but with extended metal-oxygen bond lengths. Our structural, biochemical, and computational analyses provide mechanistic insights into the ion selectivity of MgtE for Mg^2+^ over Ca^2+^.

## Introduction

Mg^2+^ ion is a fundamental biological cation implicated in various physiological functions, such as genomic stability, DNA and RNA folding, and catalysis by hundreds of enzymes^1-3^. Therefore, cellular Mg^2+^ homeostasis is vital to all domains of life and thus is strictly controlled by Mg^2+^ channels and transporters^4-7^.

MgtE is a bacterial member of the MgtE/SLC41 superfamily of Mg^2+^ channels and transporters, whose orthologs are widely conserved from bacteria to eukaryotes, including humans^8-12^. MgtE is a Mg^2+^-selective ion channel implicated in cellular Mg^2+^ homeostasis^13,14^ and is involved in bacterial survival upon exposure to antibiotics^15^. The previous single-channel recording of MgtE from *Thermus thermophilus* showed high conductance for Mg^2+^, independent of neither the pH nor the Na^+^ gradient^14,16,17^, which is consistent with the role of MgtE as a passive ion channel.

The first MgtE structure in the full-length form showed the homodimeric architecture of MgtE, where each chain consists of the transmembrane (TM) and cytoplasmic domains and a long amphipathic “plug” helix to connect these two domains^18^ (**Fig. 1a**). The MgtE cytoplasmic domain possesses regulatory Mg^2+^-binding sites to stabilize the closed state in the Mg^2+^-bound form^14,18^. In other words, the MgtE cytoplasmic domain acts as an intracellular Mg^2+^ sensor to maintain cellular Mg^2+^ homeostasis.

**Figure 1.**
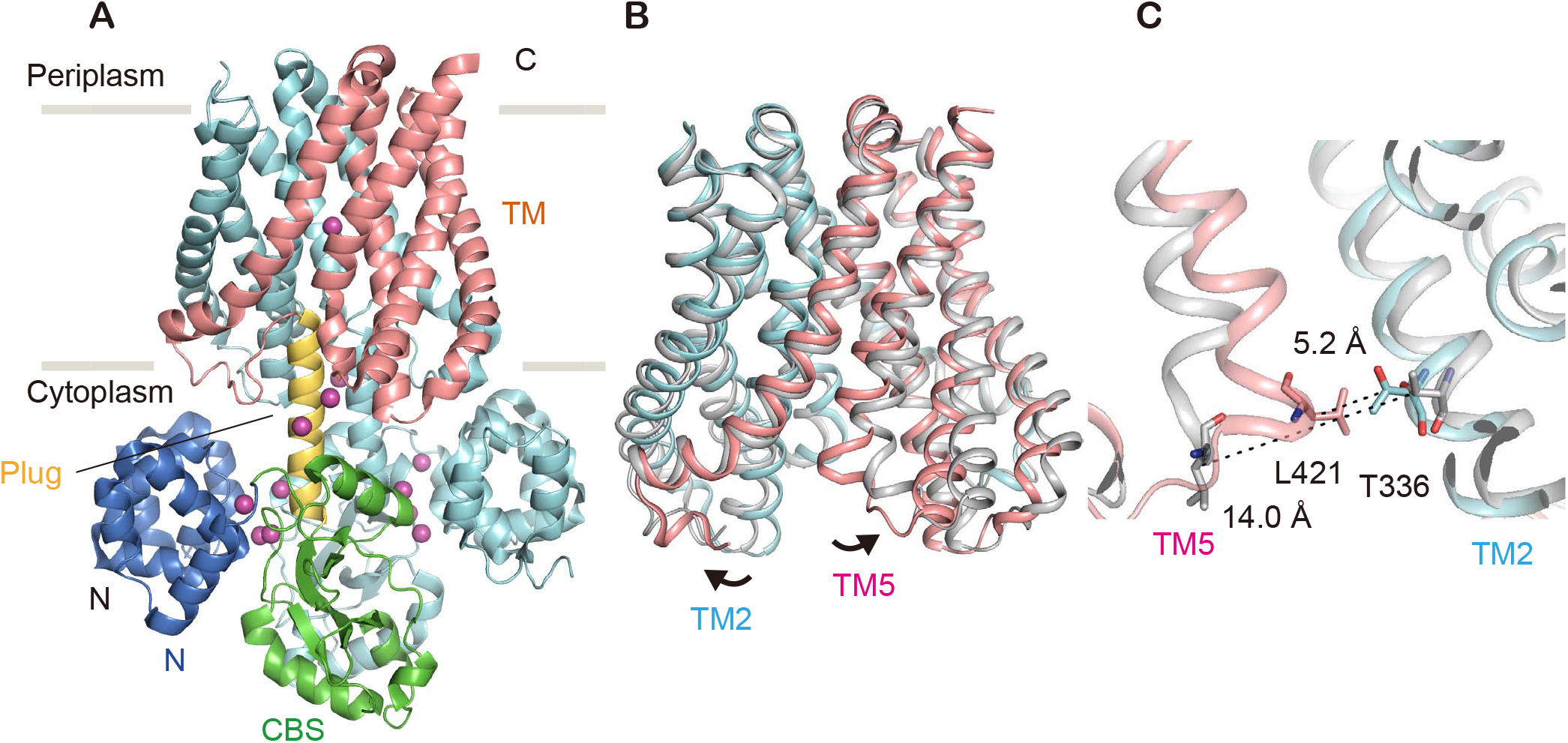
Mg^2+^-dependent cytoplasmic pore-closure motions. **(A)** *T. thermophilus* MgtE dimer structure in complex with Mg^2+^ ions (PDB ID:2ZY9), viewed parallel to the membrane. (**B, C**) The Mg^2+^-free MgtE TM domain dimer structure (PDB ID: 6LBH) superposed on the Mg^2+^-bound MgtE TM domain dimer using the Cα positions. The overall structure of the TM domain dimer (**B**) and close-up view of the dimer interface on the cytoplasmic side (**C**) are shown. The structural features of the Mg^2+^-bound structure in chain A (N, CBS and TM domains and plug helix) are colored blue, green, salmon and yellow, respectively. Chain B is colored cyan. Mg^2+^ ions are colored as magenta spheres. Mg^2+^-free MgtE is colored gray. The black arrows denote the structural transitions from the Mg^2+^-bound state to the Mg^2+^-free state (**B**). Dashed lines denote the Cα distances between Thr336 and Leu421 residues.

The subsequent crystal structures of the MgtE TM domain, in particular the one in complex with Mg^2+^ ions at high resolution (2.3 Å resolution), revealed the binding of the fully hydrated Mg^2+^ ion to the ion selectivity filter in the ion-conducting pore^16^. Nevertheless, since the previously reported MgtE TM domain structure in the presence of Ca^2+^ ions was determined only at moderate resolution (3.2 Å resolution)^16^, the mechanism of ion selectivity by MgtE, particularly the specificity for Mg^2+^ over Ca^2+^, another essential biological divalent cation, has not yet been fully understood.

Here, we determined the crystal structure of the MgtE TM domain in complex with Ca^2+^ at a higher resolution of 2.5 Å, enabling the visualization of water molecules coordinated to Ca^2+^ at the ion selectivity filter of MgtE. A combination of structural, biochemical, and computational analyses provided structural insights into the ion selectivity of MgtE for Mg^2+^ ions over Ca^2+^ ions.

## Results

### Biochemical cross-linking experiments of MgtE with Mg^2+^ and Ca^2+^

The previously reported MgtE structures in the presence and absence of Mg^2+^ ions showed Mg^2+^-dependent conformational changes in the TM domain, including the TM2 and TM5 helices, which form an ion-conducting pore (**Fig. 1b**). We verified these structural changes by cross-linking experiments with the cysteine-substituted mutant of MgtE at Leu421 and Thr336^19^, where the intersubunit distances of Cα atoms between Leu421 and Thr336 were 5.2 and 14.0 Å in the presence and absence of Mg^2+^ ions, respectively (**Fig. 1c**).

To estimate how selective the selectivity filter of MgtE in the TM domain for Mg^2+^ over Ca^2+^ is, using the MgtE T336C/L421C mutant lacking the N domain (MgtE ΔN T336C/L421C) and Cu^2+^ phenanthroline as a catalyst, we performed biochemical cross-linking experiments in the presence of Mg^2+^ or Ca^2+^ at a concentration of gradients (**Fig. 2**). Whereas MgtE also possesses regulatory Mg^2+^ binding sites in the cytoplasmic domain, deletion of the N domain is known to abolish the Mg^2+^-sensitivity of the cytoplasmic domain^14^. Therefore, in this cross-linking experiment, to estimate the Mg^2+^/Ca^2+^ affinity to the ion selectivity filter in the MgtE TM domain as exactly as possible, we employed the ΔN mutant to exclude the influence of Mg^2+^ binding to the cytoplasmic domain.

**Figure 2.**
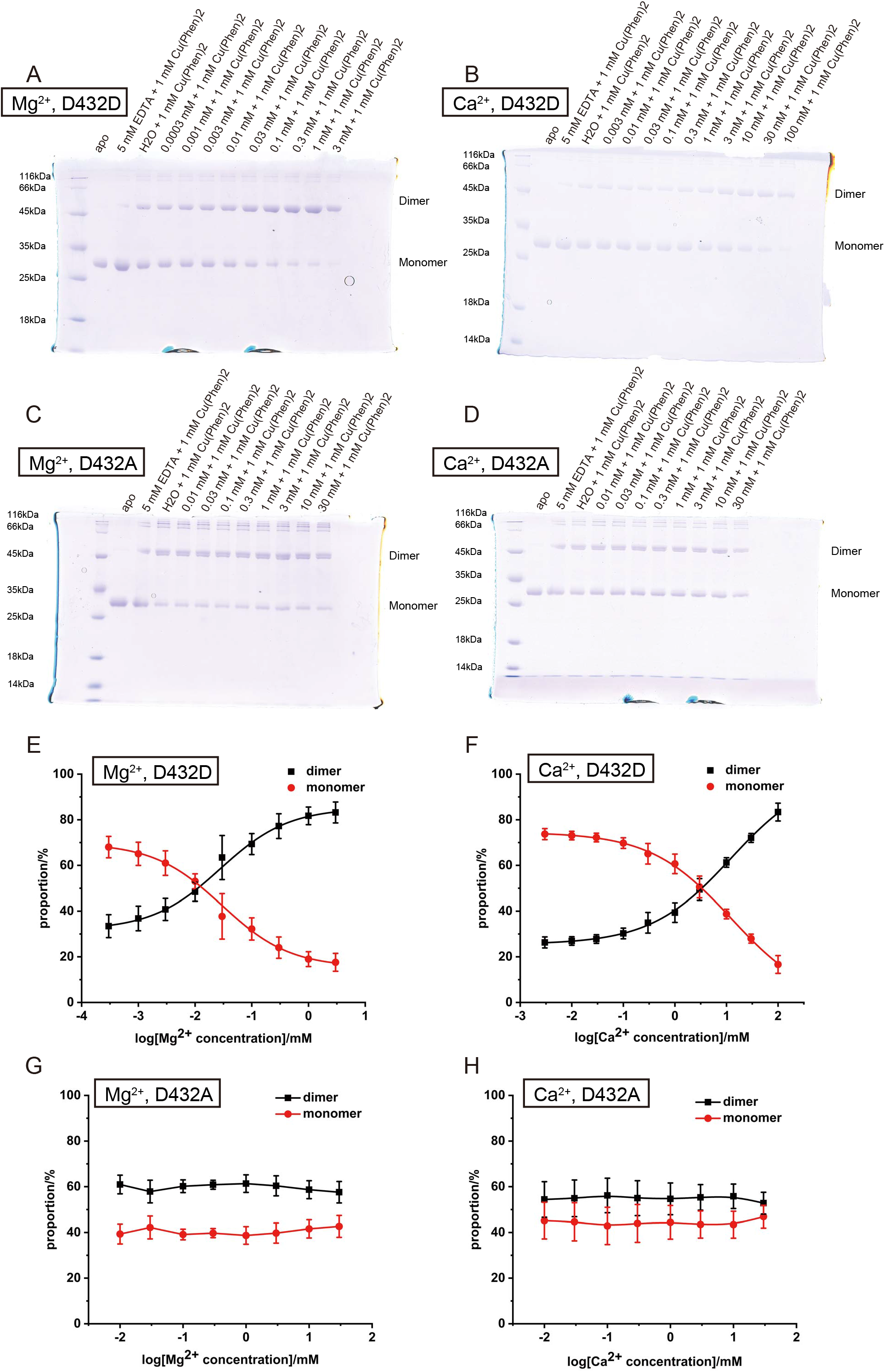
Biochemical cross-linking experiments of MgtE with Mg^2+^ and Ca^2+^. **(A-D)** Representative SDS–PAGE gels from biochemical cross-linking experiments of the MgtE ΔN domain double cysteine mutant T336C/L421C with Mg^2+^ (**A**), Ca^2+^ (**B**), Mg^2+^ and the D432A mutation (**C**), and Ca^2+^ and the D432A mutation (**D**). (**E-H**) Densitometric quantification of SDS–PAGE band intensities of the MgtE dimer (black) and monomer (red) with Mg^2+^ (**E**), Ca^2+^ (**F**), Mg^2+^ and the D432A mutation (**G**) and of Ca^2+^ and the D432A mutation (**H**) for sigmoidal curve fitting. Experiments were performed six times. Error bars represent SEM. All SDS–PAGE gels can be found in **Supplementary Figs. 1-4**.

As the concentrations of Mg^2+^ and Ca^2+^ increased, the MgtE ΔN T336C/L421C mutant exhibited a stronger band corresponding to the MgtE dimer (**Fig. 2**). The estimated EC_50_ values of 27.7±3.7 μM and 12.2±1.9 mM for Mg^2+^ and Ca^2+^, respectively, indicated the high selectivity of Mg^2+^ over Ca^2+^. Notably, the alanine substitution of Asp432 in the TM5 helix, which forms the ion selectivity filter called the M1 site (**Fig. 3a**), abolished the divalent cation-dependent cross-linking (**Fig. 2**), indicating that the divalent cation binding to the M1 site indeed induces chemical cross-linking. Overall, these results indicate that the ion selectivity filter of MgtE is highly selective for Mg^2+^ over Ca^2+^.

**Figure 3.**
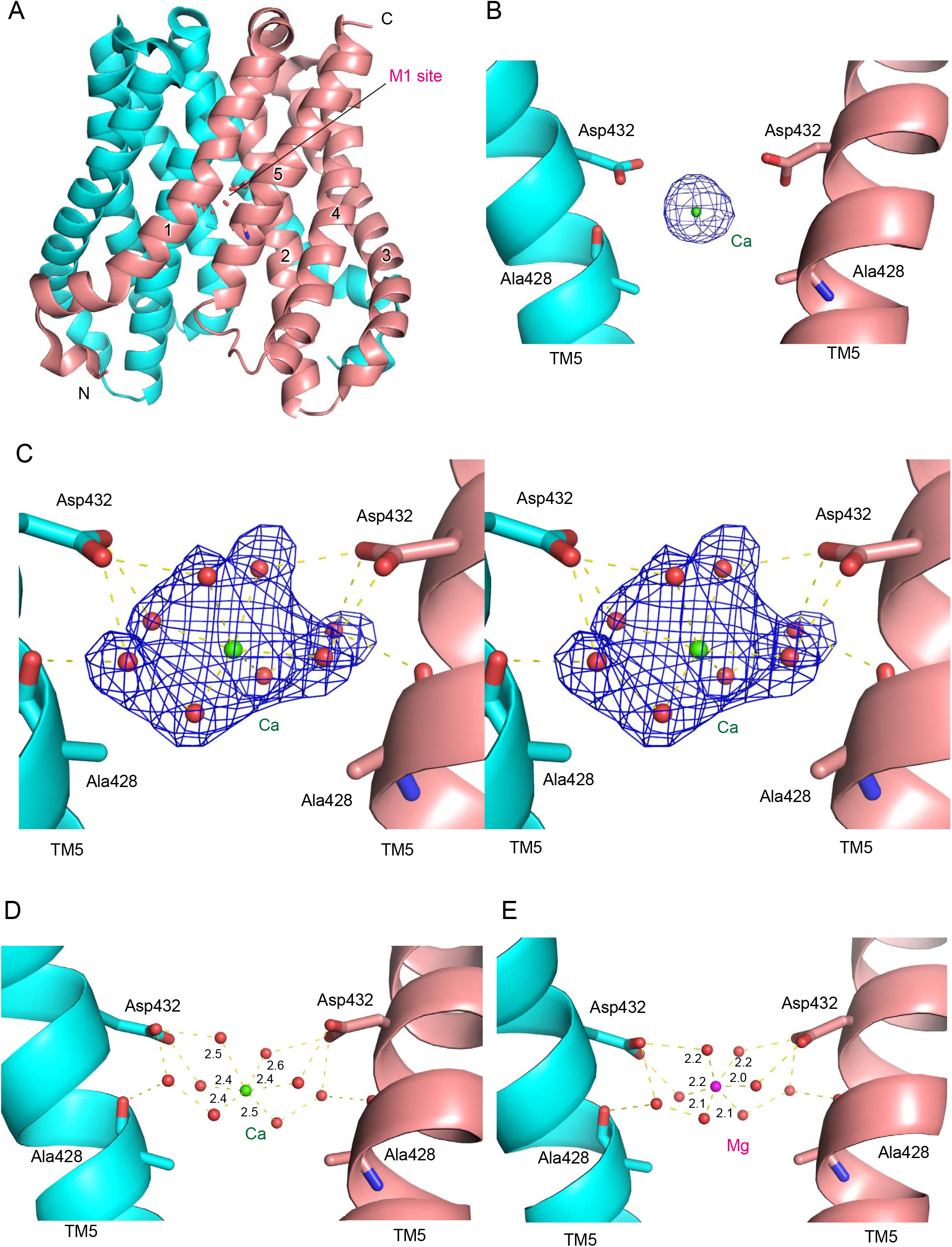
Ion selectivity filter. **(A)** The Mg^2+^-bound MgtE TM domain structure (PDB ID: 4U9L). The location of the ion selectivity filter (M1 site) is marked. (**B, C, D**) A close-up view of the M1 site in the Ca^2+^-bound MgtE TM domain structures, previously determined at 3.2 Å resolution (PDB ID: 4WIB) (**B**) and determined at 2.5 Å resolution in this study in a stereo view. (**C, D**). (**E**) A close-up view of the M1 site in the Mg^2+^-bound MgtE TM domain structure (PDB ID: 4U9L). The POLDER-OMIT maps for Ca^2+^ and associated water molecules are shown in blue mesh (contoured at 3.0 σ) (**B, C**). The coloring scheme is the same as in **Fig. 1**. Amino acid residues at the metal binding site are shown in stick representation. Mg^2+^, Ca^2+^ and water molecules are shown as magenta, green and red spheres, respectively. Dashed lines indicate hydrogen bonds, and associated numbers show the water-metal distances (Å).

### Higher-resolution crystal structure of MgtE in the Ca^2+^-bound form

To obtain a higher-resolution structure of MgtE in complex with Ca^2+^ ions, we crystallized the MgtE TM domain in the presence of Ca^2+^ ions using the lipidic cubic phase (LCP) technique^20^, collected the datasets from a number of microcrystals using the ZOO automated data collection system^21^, and merged the datasets from 209 microcrystals using the KAMO automated data processing system^22^. The final datasets yielded a resolution of 2.5 Å, higher than before (3.2 Å). The newly determined structure of the MgtE TM domain is essentially the same as the previously determined MgtE TM domain (PDB ID: 4U9L), with an RMSD of 0.34 Å for Cα atoms.

In this structure, we observed more detailed features in the electron density maps at the ion selectivity filter (**Fig. 3c**) than could be observed in the previous, lower-resolution Ca^2+^-bound structure (**Fig. 3b**). Among the possible coordination geometries of Ca^2+^ ions in biology, with a range of water coordination numbers from six to eight^23-25^, the electron densities in our structure adopt an octahedral coordination geometry of Ca^2+^ ions with six water molecules in the first hydration shell (**Fig. 3c**). The side chains of Asp432 residues interact with four of six water molecules in the first hydration shell (**Fig. 3d**). In addition, two extra water molecules in the second hydration shell form hydrogen bonds with the side chain of Asp432 residues and main chain carbonyl oxygen atoms of Ala428 residues (**Fig. 3d**). These interactions seemingly stabilize Ca^2+^ ions in the M1 site in the fully hydrated form. The bonding distances between Ca^2+^ ions and coordinated water molecules are 2.4–2.6 Å (**Fig 3D**), which is consistent with the range of Ca-O distances (2.2-2.7 Å) in previously reported crystal structures^3^.

Notably, in the previously reported Mg^2+^-bound structure (**Fig. 3e**), Mg^2+^ also adopts a very similar octahedral coordination geometry to that observed in the present Ca^2+^-bound structure (**Fig. 3d**) but with shorter bonding distances between Mg^2+^ and water molecules of 2.0–2.2 Å, which is also consistent with the typical Mg-O distance in previously reported crystal structures^3^.

### MD simulations

To further examine Mg^2+^ and Ca^2+^ recognition by the M1 site of MgtE, we performed MD simulations based on the current Ca^2+^-bound crystal structure together with the previously reported Mg^2+^-bound structure (**Fig. 4**). The overall structures were mostly stable during the 1-μs simulations starting from the MgtE structure embedded in the POPC lipid bilayer (**Fig. 4a**). Both the Mg^2+^ ion and Ca^2+^ ion were stably bound to the M1 site with all six water molecules in the fully hydrated state. Importantly, the distances between Mg^2+^ and water molecules and between Ca^2+^ and water molecules were stable during the simulations, whereas Ca^2+^ maintained a longer distance from water molecules than Mg^2+^ (**Fig. 4b, c, d**).

**Figure 4.**
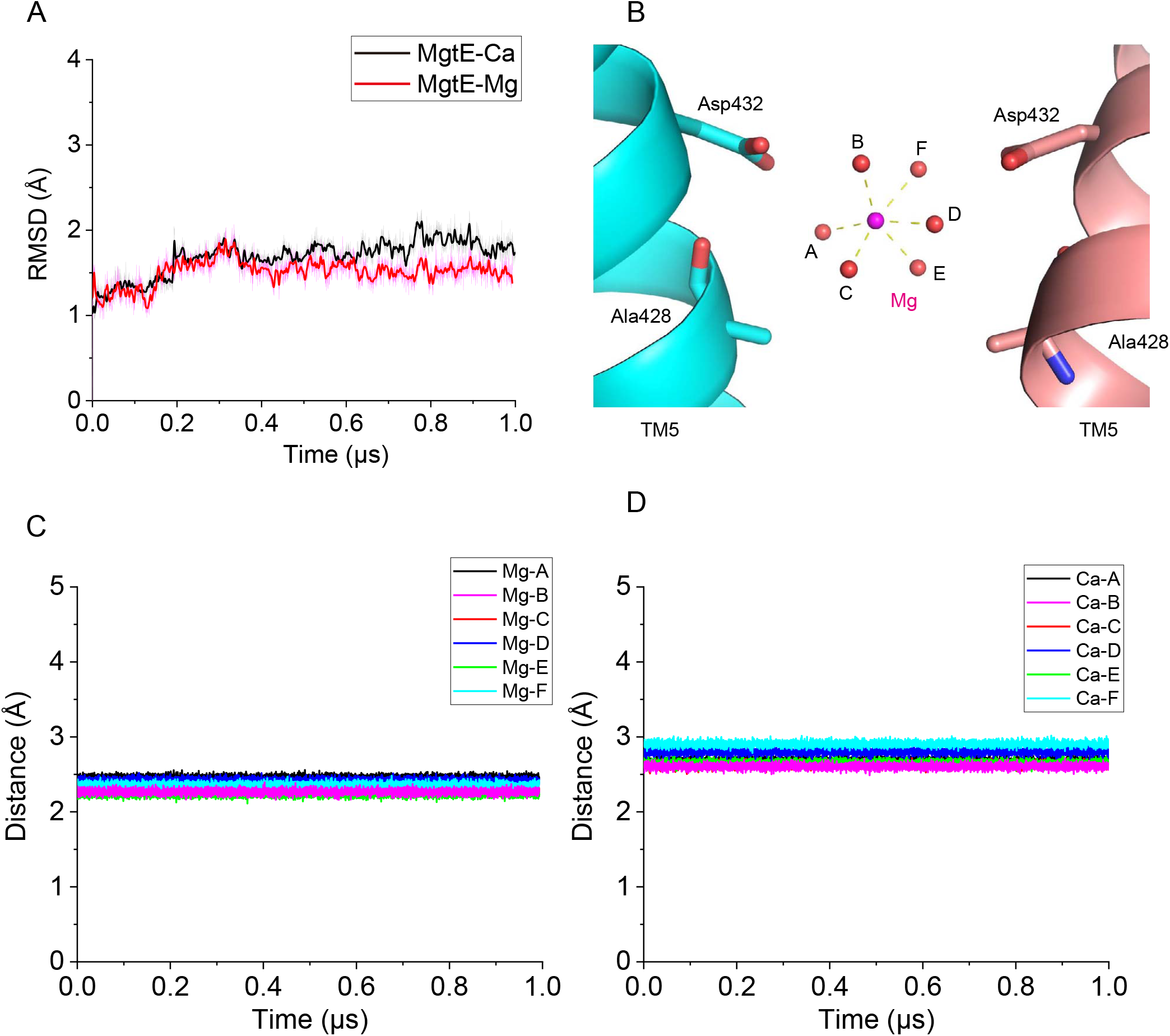
MD simulations. **(A)** Structural deviations from the MgtE TM domain structure during 1-μs MD simulations with Mg^2+^ or Ca^2+^ ions. (**B**) A close-up view of the metal binding site in the MgtE TM domain structure with the fully hydrated Mg^2+^ ion (PDB ID: 4U9L). Alphabetical labels of the water molecules correspond to those in Fig. 4c, d. (**C, D**) Water-metal distances between Mg^2+^ and coordinated water molecules (**C**) and between Ca^2+^ and coordinated water molecules (**D**) during 1-μs -MD simulations.

These results further support the insights from the crystal structures that the M1 site of MgtE can accommodate both Mg^2+^ and Ca^2+^ in the octahedral coordination geometry with six water molecules but with longer bonding distances between Ca^2+^ and water molecules.

## Discussion

Mg^2+^ and Ca^2+^ are fundamental divalent cations for life. However, relatively little is known about the selectivity mechanisms by which ion channels and transporters discriminate them. In this work, we showed by biochemical cross-linking that the M1 site of the MgtE TM domain is highly selective for Mg^2+^ over Ca^2+^ (**Fig. 2**). The improved crystal structure of the MgtE TM domain in complex with Ca^2+^ together with the MD simulations suggested that the M1 site recognizes Ca^2+^ in an octahedral coordination geometry with six water molecules, similar to that observed in the previously determined Mg^2+^-bound structure, but with longer metal-water bond lengths (**Figs. 3 and 4**).

Based on these results, we discuss the ion selectivity mechanism of MgtE for Mg^2+^ over Ca^2+^. First, the coordination number can be from six to eight for Ca^2+^, but seven is most common in aqueous solution^23-26^. Consistently, the Ca^2+^ ATPase pump and Na^+^/Ca^2+^ exchanger also recognize Ca^2+^ ions with a coordination number of seven^27,28^, which directly explains their ion selectivity for Ca^2+^ over Mg^2+^, since Mg^2+^ ions have a strict octahedral coordination with six water molecules. On the other hand, in the case of MgtE, the M1 site recognizes Ca^2+^ ions in the octahedral coordination geometry with six water molecules (**Fig. 3**). Since Ca^2+^ is estimated to have a coordination number from six to eight and since the coordination number of seven is preferable, a loss of coordination would occur when Ca^2+^ takes on an octahedral coordination number in the M1 site of MgtE. In other words. Ca^2+^ has to be forced into a lower coordination mode when bound to the M1 site of MgtE. This may explain the selectivity of MgtE for Mg^2+^ over Ca^2+^.

Overall, our structural, biochemical and computational analyses provide insights into the selectivity of MgtE for Mg^2+^ over Ca^2+^.

## Methods

### Expression and purification

The MgtE ΔN domain mutant gene from *T. thermophilus* (residues 130-450) was subcloned into a pET28a vector containing an N-terminal hexahistidine tag and a thrombin cleavage site. The human rhinovirus 3C (HRV3C) protease cleavage site was inserted between residues Asp267 and Val268 at the loop region between the cytoplasmic domain and TM domain. The protein expression and purification of MgtE were similarly performed as described previously^16,18,19^. MgtE protein was overexpressed in *E. coli* Rosetta (DE3) cells in LB medium containing 30 μg/ml kanamycin at 37 °C by adding 0.5 mM isopropyl-β-D-thiogalactoside (IPTG) at an OD_600_ of ∼0.5, and then *E. coli* cells were further cultured at 18 °C for 16 hours. The *E. coli* cells were harvested by centrifugation (6,000 × g, 15 minutes) and then resuspended in buffer H [150 mM NaCl, 50 mM HEPES (pH 7.0) and 0.5 mM phenylmethanesulfonyl fluoride (PMSF)]. All purification procedures were performed at 4 °C. The *E. coli* cells were disrupted with a microfluidizer. After centrifugation (20,000 × g, 30 minutes), the supernatants were collected and subjected to ultracentrifugation (200,000 × g, 1 hour). The membrane fraction from ultracentrifugation was then solubilized with buffer S [300 mM NaCl, 50 mM HEPES (pH 7.0), 2% (w/v) n-dodecyl-beta-d-maltopyranoside (DDM) (Anatrace, USA) and 0.5 mM PMSF] for 2 hours. The solubilization fraction was loaded onto a Ni-NTA column preequilibrated with buffer A [300 mM NaCl, 50 mM HEPES (pH 7.0) and 0.05% (w/v) DDM] containing 20 mM imidazole, mixed, and incubated for 1 hour. The Ni-NTA column was washed with buffer A containing 50 mM imidazole, and the MgtE protein was eluted with buffer A containing 300 mM imidazole.

To cleave the HRV3C protease cleavage site, the eluate was mixed with Ni-NTA beads preequilibrated in buffer B and His-tagged HRV3C protease and then dialyzed against buffer B overnight. The sample was reloaded on a column, and the flow-through fractions containing the MgtE TM domain protein were concentrated using an Amicon Ultra 50 K filter (Merck Millipore, USA). After concentration, the sample was injected into a Superdex 200 10/300 size-exclusion column (GE Healthcare, USA) equilibrated with buffer C [25 mM HEPES (pH 7.0), 150 mM NaCl and 0.025% (w/v) DDM] for size-exclusion chromatography (SEC). The peak fractions containing the MgtE TM domain protein were collected and concentrated to 10 mg/ml using an Amicon Ultra 50 K filter (Merck Millipore, USA) for crystallization.

For the MgtE ΔN T336C/L421C and MgtE ΔN T336C/L421C/D432A cysteine mutants, the protein expression and preparation of the membrane fractions were performed similarly to the methods described above. The membrane fractions were solubilized with buffer D [50 mM HEPES (pH 7.0), 150 mM NaCl, 2% DDM, 20 mM imidazole, 1 mM PMSF, 1 mM β-mercaptoethanol (β-ME)] for 2 hours. Then, insoluble materials were removed by ultracentrifugation (200,000 × g, 1 hour). The supernatant was mixed with Ni-NTA resin preequilibrated with buffer D, incubated for 1 hour, washed with buffer E [50 mM HEPES (pH 7.0), 150 mM NaCl, 0.05% DDM, 50 mM imidazole, 1 mM β-ME], and then eluted with buffer F [50 mM HEPES (pH 7.0), 150 mM NaCl, 0.05% DDM, 300 mM imidazole, 1 mM β-ME]. The eluted MgtE proteins were dialyzed in buffer G [50 mM HEPES (pH 7.0), 150 mM NaCl, 0.05% DDM, 20 mM dithiothreitol (DTT)] overnight and applied to a Superdex 200 10/300 size-exclusion column equilibrated with buffer H [20 mM HEPES (pH 7.0), 150 mM NaCl, 0.03% DDM] for SEC. The peak fractions were concentrated to 0.5 mg/ml using an Amicon Ultra 50K filter.

### Crystallization

Before crystallization, the purified MgtE TM domain protein was mixed with CaCl_2_ at a final concentration of 100 mM and incubated on ice for 30 minutes. The protein was then mixed with monoolein (NU-CHEK, USA) at a ratio of 2:3 (w:w) in a twin syringe to generate lipidic cubic phase (LCP)^20^. For crystallization, a Gryphon LCP crystallization robot (Art Robbins Instruments, USA) was employed to dispense 50 nl of LCP drops onto a 96-well sandwich plate and to overlay 700 nl reservoir solutions. Crystals appeared at 18 °C after one week in the reservoir solution containing 30% (w/v) polyethylene glycol (PEG) 400, 100 mM HEPES (pH 7.5), and 100 mM NaSCN.

### X-ray data collection and structure determination

X-ray diffraction data were collected at the BL32XU beamline at SPring-8 (Harima, Japan) using the ZOO automatic data collection system^21^ and processed with KAMO^22^ and XDS^29^. The structure of the MgtE TM domain in complex with Ca^2+^ (residues 271-448 for chains A and B) was determined by molecular replacement with Phaser^30^ using the Mg^2+^-bound MgtE TM domain structure (PDB ID: 4U9L). The atomic model was then manually built using COOT^31^ and refined with PHENIX^32^. The Ramachandran plots were calculated using MolProbity^33^. X-ray data collection and refinement statistics are summarized in Table 1. All structure figures were generated using PyMOL (https://pymol.org/).

**Table 1.**
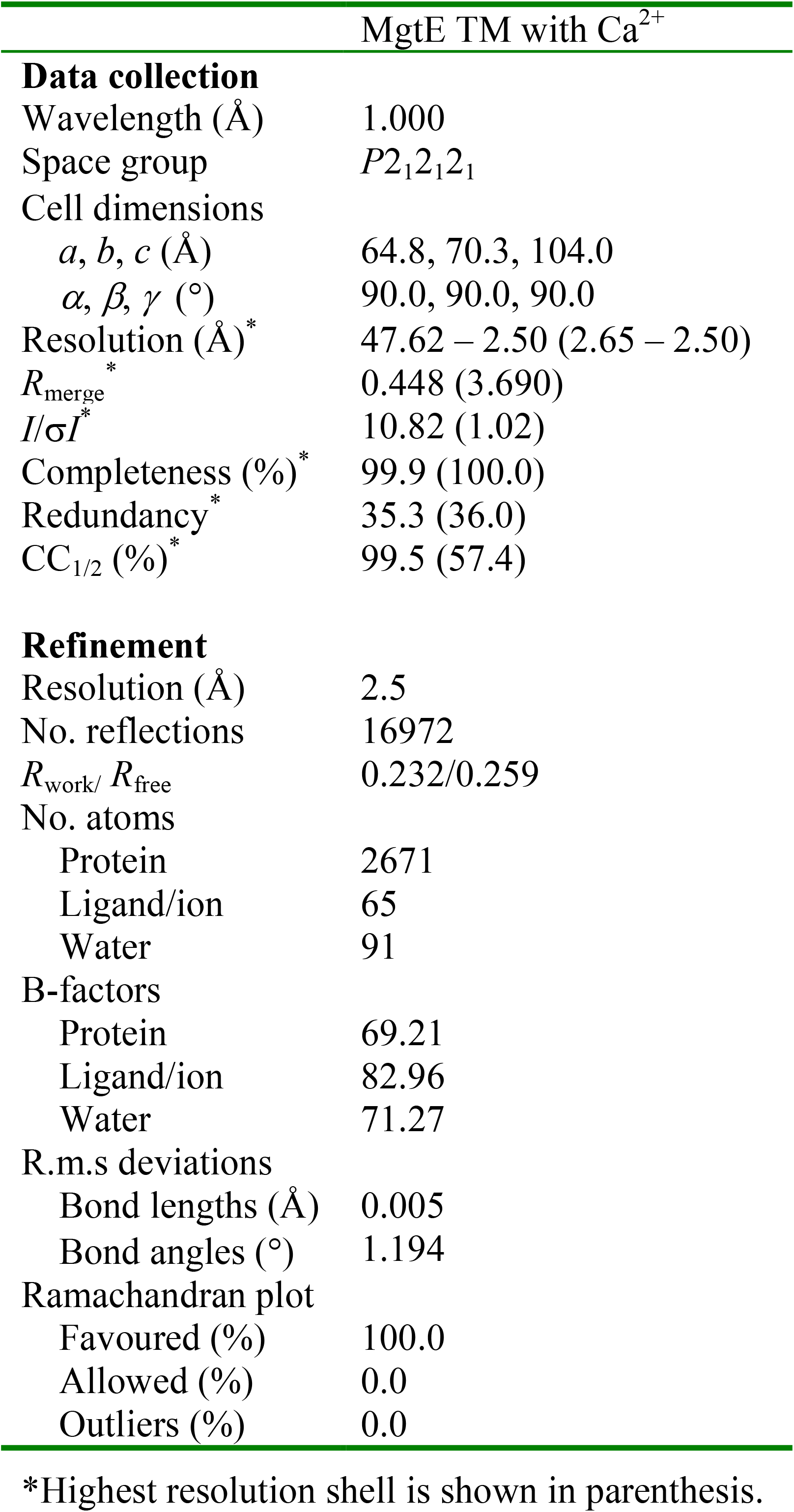
X-ray data collection and refinement statistics.

### Biochemical cross-linking

Biochemical cross-linking experiments were performed as described previously^19^. First, 4 μl of MgtE protein at 0.5 mg/ml was mixed with 0.5 μl of EDTA at a final concentration of 5 mM, MgCl_2_ or CaCl_2_, at appropriate concentrations, respectively, incubated on ice for 30 min, then mixed with 0.5 μL of 10 mM freshly prepared Cu^2+^ bis-1,10-phenanthroline (with molar ratio 1:3) to react on ice for another 30 min. Samples were analyzed by SDS–PAGE in nonreducing conditions. Experiments were repeated six times. SDS–PAGE gels were quantified by ImageJ (NIH, USA), and the quantified data were fitted to a nonlinear curve by Origin (OriginLab, USA).

### Molecular dynamics simulations

Molecular dynamics (MD) simulations were performed using ‘Desmond’ software^34^. The initial positioning of MgtE in the 1-palmitoyl-2-oleoyl-sn-glycero-3-phosphocholine (POPC) membrane was obtained from the OPM database^35^. The POPC membrane-bound structures were built using a simple point charged water model (SPC) in an orthorhombic box with dimensions of 10 Å ×10 Å ×10 Å. To maintain balance and neutralize the system, counter ions (Na^+^ or Cl^−^) were added. NaCl (150 mM) was added to the simulation box to represent background salt under physiological conditions. Prior to MD simulation, the DESMOND default relaxation protocol was applied to each system. (1) 100 ps simulations in the NVT ensemble with Brownian kinetics using a temperature of 10 K with solute heavy atoms constrained; (2) 12 ps simulations in the NVT ensemble using a Berendsen thermostat with a temperature of 10 K and small-time steps with solute heavy atoms constrained; (3) 12 ps simulations in the NPT ensemble using a Berendsen thermostat and barostat for 12 ps simulations at 10 K and 1 atm, with solute heavy atoms constrained; (4) 12 ps simulations in the NPT ensemble using a Berendsen thermostat and barostat at 300 K and 1 atm, with solute heavy atoms constrained; (5) 24 ps simulations in the NPT ensemble using a Berendsen thermostat and barostat at 300 K and 1 atm without constraint. After equilibration, the MD simulations were performed for 1000 ns. Long-range electrostatic interactions were computed using a smooth particle mesh Ewald method. The trajectory recording interval was set to 200 ps, and other default parameters of DESMOND were used during MD simulation runs. All simulations used the all-atom OPLS_2005 force field^36,37^, which was used for the protein ions, and ligand molecules, for proteins, ions, lipids and SPC waters. All simulations were run on a DELL T7920 graphic working station (with an NVIDA Tesla K40C-GPU). Analysis and visualization were performed on a 12-CPU CORE DELL T3610 graphic working station.

### Statistics and reproducibility

The biochemical cross-linking experiments in **Fig. 2** were repeated six times. Error bars represent the standard error of the mean. X-ray data collection and refinement statistics are summarized in **Table 1**.

## Data availability

The atomic coordinates and structure factors of MgtE were deposited in the Protein Data Bank (PDB ID: 7F7U). All SDS–PAGE gels can be found in **Supplementary Figs. 1-4**. All materials are available from the authors upon reasonable request.

## Acknowledgments

We thank the staff from the BL32XU and BL41XU beamlines at SPring-8 and from BL17U1 at Shanghai Synchrotron Radiation Facility (SSRF) for assistance during X-ray data collection. The diffraction experiments were performed at SPring-8 BL32XU and BL41XU (Proposal Nos. 2019A2514 and 2020A2524) and at SSRF BL17U1 (Proposal No. 2018-SSRF-PT-004257). This work was supported by funding from the Ministry of Science and Technology of China (National Key R&D Program of China: 2016YFA0502800) to M.H. and from the National Natural Science Foundation of China (32071234) to M.H. This work was also supported by the Innovative Research Team of High-level Local Universities in Shanghai and a key laboratory program of the Education Commission of Shanghai Municipality (ZDSYS14005) and by the Open Research Fund of State Key Laboratory of Genetic Engineering, Fudan University (No. SKLGE-2105).

## Author contributions

X.T. and D.S. purified and crystallized the MgtE TM domain and determined the Ca^2+^-bound structure of MgtE. X.T. performed the biochemical cross-linking experiments. J.W. and Y.Y. performed MD simulations. X.T., D.S. and M.H. wrote the manuscript. M.H. supervised the research. All authors discussed the manuscript.

