## Supplementary Figure 1-4 for "Structural insights into the ion selectivity of the MgtE channel for Mg^2+^ over Ca^2+^"

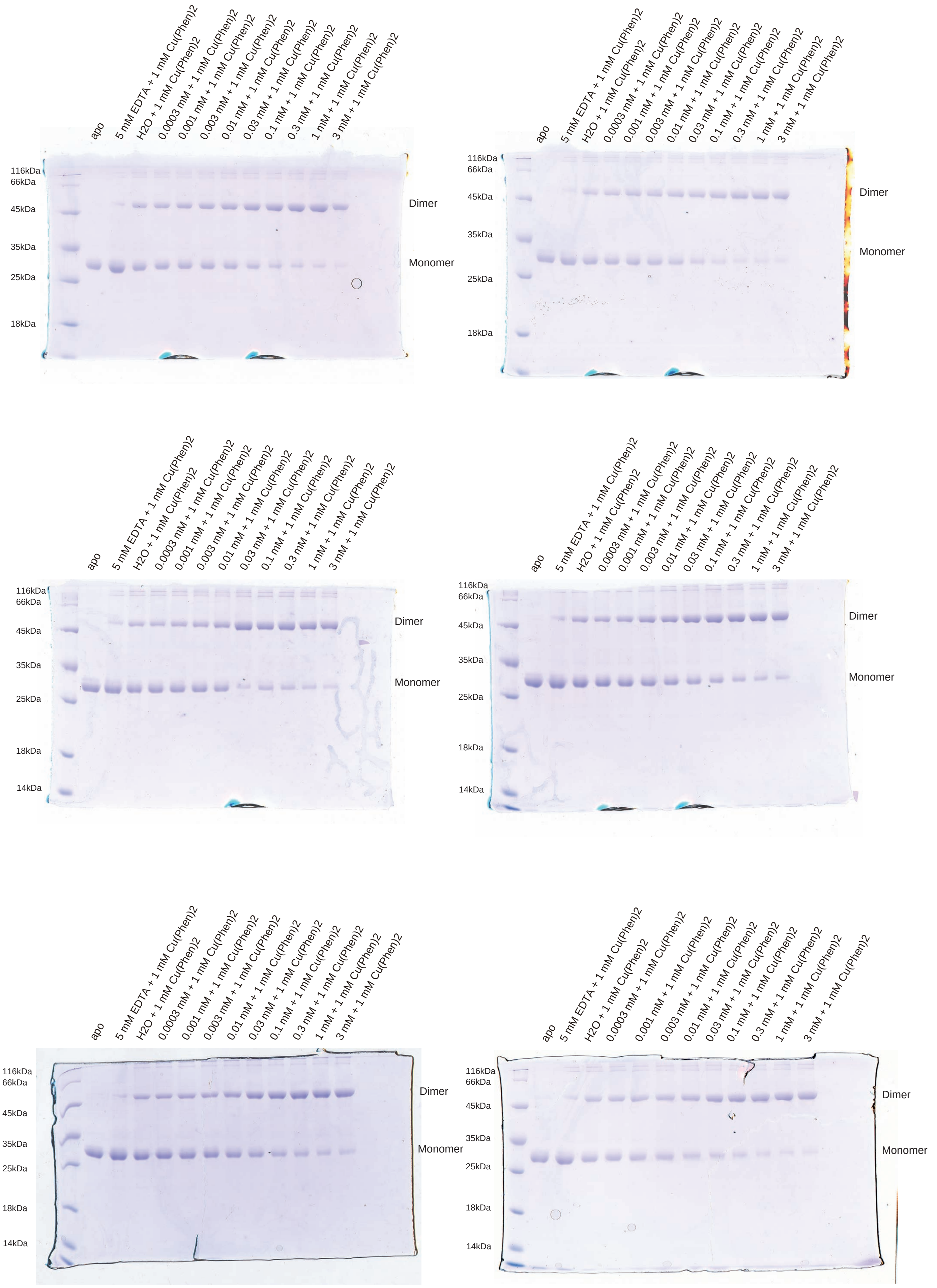


**Supplementary Fig. 1. SDS–PAGE gels from biochemical cross-linking experiments with MgtE ∆N T336C/L421C and $Mg^{2+}$.**

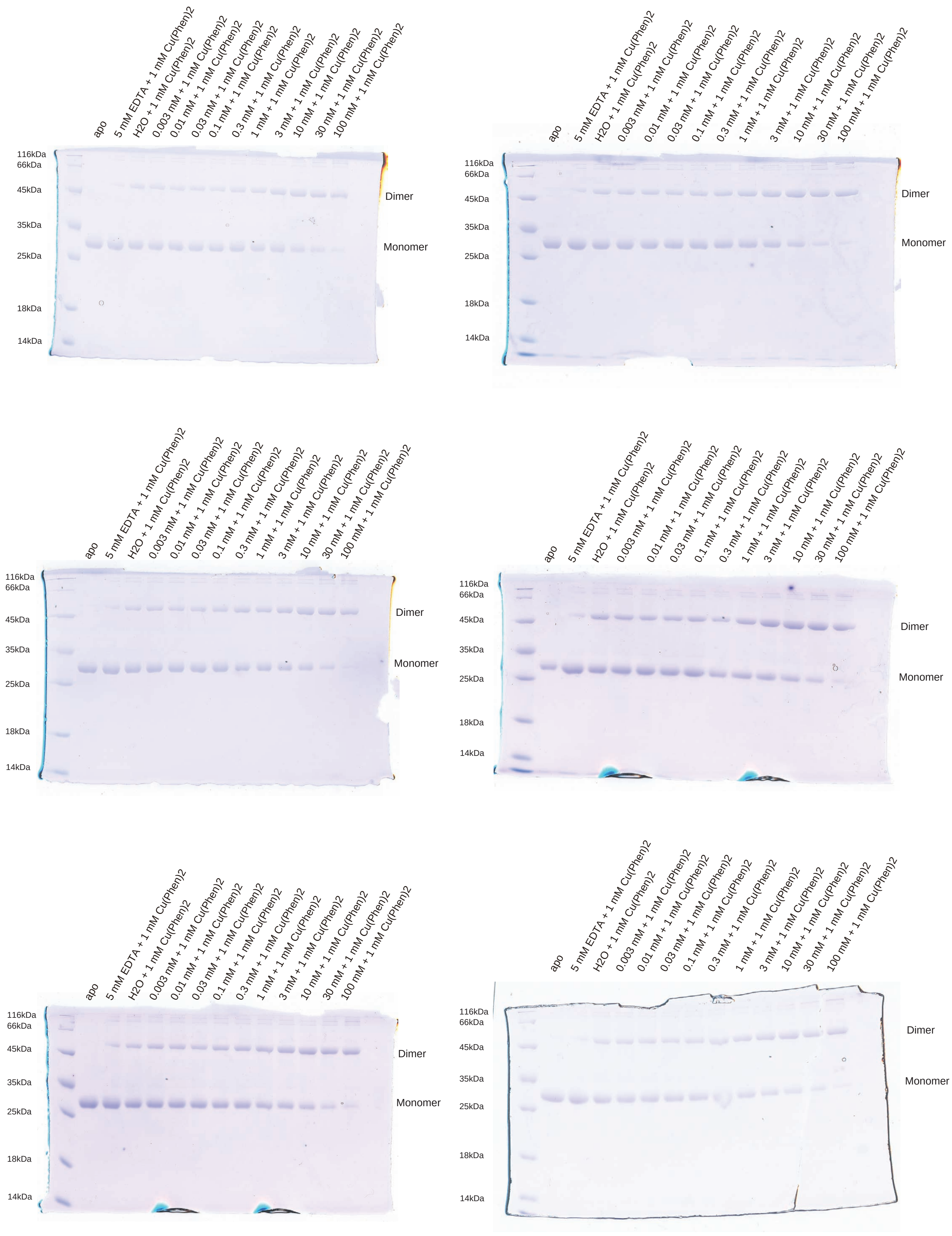


**Supplementary Fig. 2. SDS–PAGE gels from biochemical cross-linking experiments with MgtE ΔN T336C/L421C and $Ca^{2+}$.**

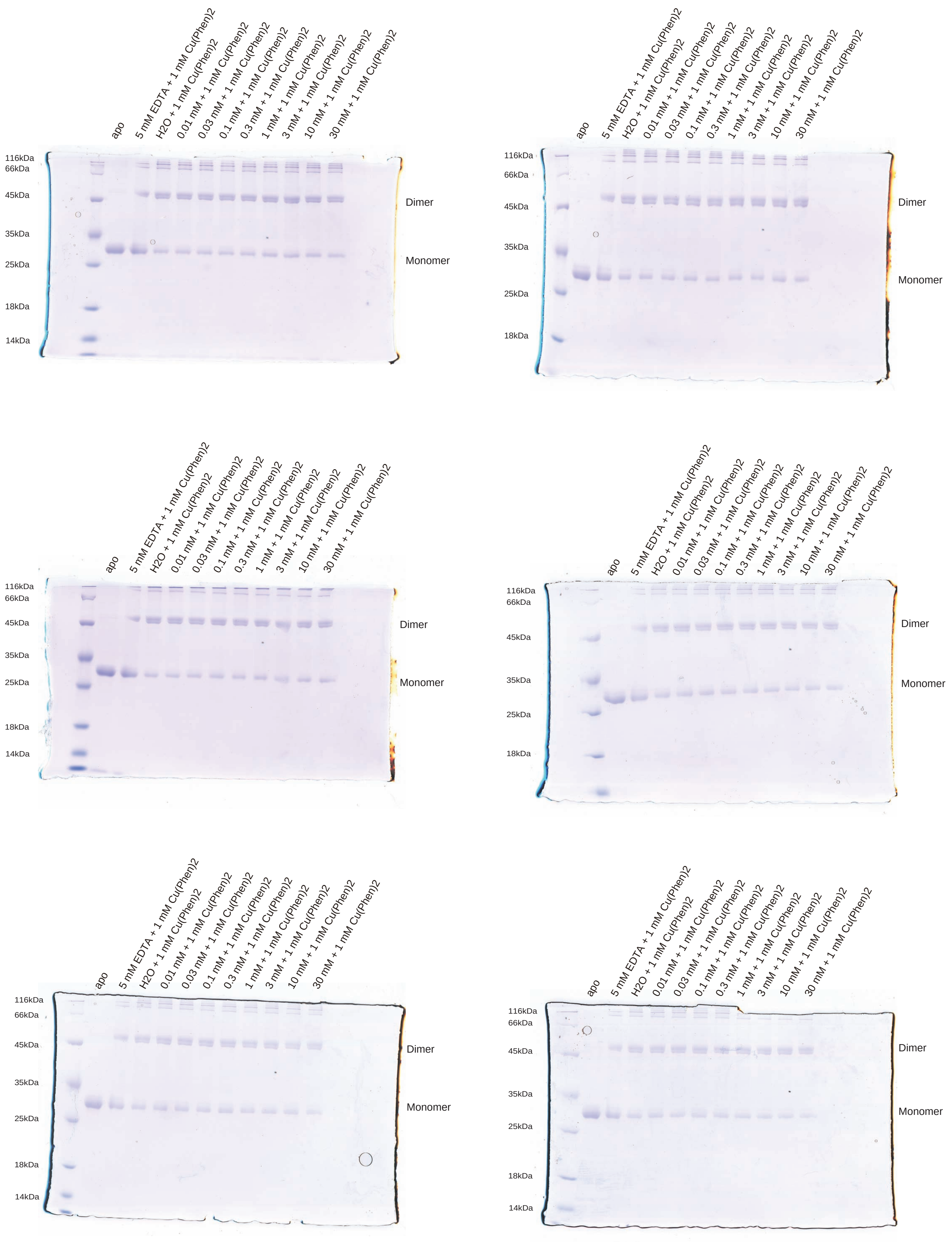


**Supplementary Fig. 3. SDS–PAGE gels from biochemical cross-linking experiments with MgtE $\Delta$N T336C/L421C/D432A and $Mg^{2+}$.**

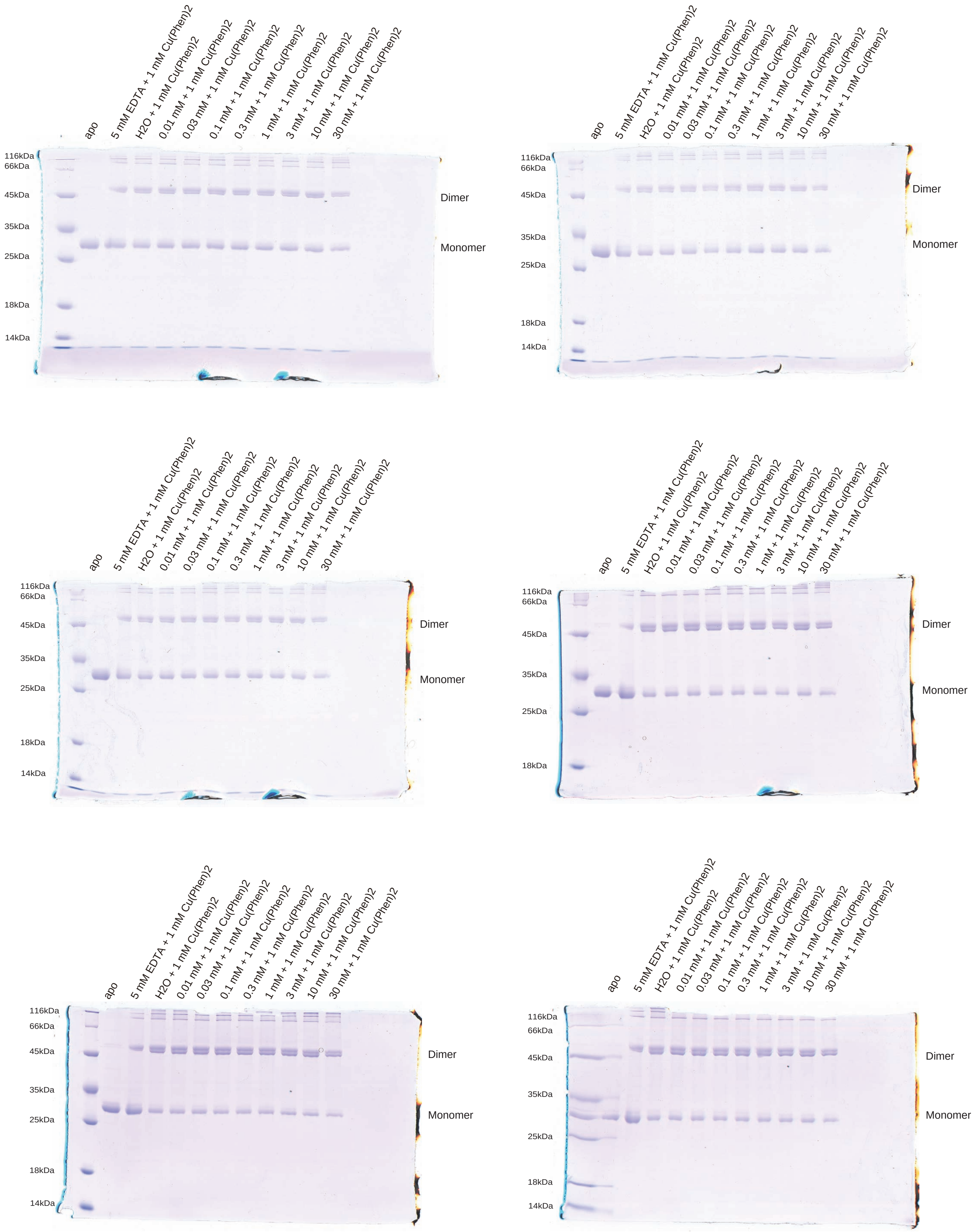


**Supplementary Fig. 4. SDS–PAGE gels from biochemical cross-linking experiments with MgtE ΔN T336C/L421C/D432A and $Ca^{2+}$.**
